# A deep learning phenotyping method for genetic analysis of 3D micro-CT data

**DOI:** 10.1101/2023.08.24.554725

**Authors:** A. Karshenas, T. Linderoth, R. Zatha, B. Rusuwa, R. Durbin

## Abstract

The number of Genome-Wide Association Studies (GWAS) has been growing rapidly in recent years due to developments in genotyping and sequencing platforms. When applied to quantitative traits, these and other statistical genetics approaches require large amounts of consistently and accurately measured phenotypes. Here, we introduce a computational toolbox based on deep convolutional neural networks that we have developed to phenotype quantitative traits describing morphology from micro-CT-scan image datasets. We illustrate the use of this Deep Learning Phenotyper (DLP) on a sample set of craniofacial CT scans of 118 samples from two very closely related species of Lake Malawi cichlid fish, *Maylandia zebra* and *Cynotilapia zebroides*. We show that the pipeline constructed and implemented here is capable of measuring morphological skeletal phenotypes with high accuracy. We also demonstrate how this pipeline can be integrated with existing GWAS frameworks to identify candidate association loci. We believe the methods we present here will be valuable for groups studying quantitative morphological traits not only in fishes, but in other vertebrates using CT scan datasets.

## 1 Introduction

One of the most striking but complex ways that individuals vary within and between species is in their three-dimensional shape, or morphology. The genetic basis of morphological variation has been studied in many vertebrates beyond humans in the last few decades (Roff and Mousseau, 1987; Wallace et al., 2014). Although landmarking and other manual or semi-manual methods exist for quantification of morphology using live samples, photographs, and three-dimensional computer tomographic (CT) images, they are hard to scale to the large datasets that are required for statistical genetic analyses (Webster and Sheets, 2010). In this work, we introduce a deep learning-based, three-dimensional phenotyping method that can be used to scale up and automate morphometric measurements from CT scan data.

Since the development of tomography methods and the first CT scan of a living human brain in 1971 (Hounsfield, 1973), CT scanning instruments have become globally available, and are widely used in medical diagnosis, rapid use of CT scans in COVID-19 diagnosis being a perfect example (Li and Xia, 2020), as well as for other purposes. In particular, they have been used to generate high-resolution images of skeletons and soft tissues in many vertebrates for morphometric research (Steger et al., 2012; Khmelinskii et al., 2012). Raw CT images are generated through a rotating pair of conventional X-ray generators and detectors, that provide a stack of x-ray projections captured from different orientations. These images generated by the scanners then go through tomography algorithms in order to generate CT images (Withers et al., 2021). These methods back calculate the density of materials based on x-ray absorption (Hounsfield, 1980), resulting in high-resolution three-dimensional images that are commonly stored as stacks of two-dimensional (2D) images, typically ordered along the anterior-posterior axis of the body, giving radiologists and researchers access to detailed images of internal organs and skeleton structure.

These CT methods have started to generate large datasets across samples that can be used with genetic data to uncover causal links in phenotypic variation observed both within and across species. Classic examples of such studies are Genome-Wide Association Studies (GWAS) (Frary et al., 2000; Burton et al., 2007; Wood et al., 2014; He et al., 2017; Buniello et al., 2018; Soliai et al., 2021) and heritability analysis (Visscher et al., 2008). While genetic analysis of simple morphological traits (such as human height) is routine (Wood et al., 2014), there is a desire to apply the GWAS technology to more complex multi-dimensional phenotypes. The power of these computational approaches increase with the sample size (Spencer et al., 2009; Hong Eun Pyo, 2012), making accurate, high-throughput phenotyping pipeline desirable. Hence, our aim was to develop an unbiased, reliable, and easy-to-use toolbox that can be applied, with minor adjustments, to CT scans of most vertebrates that is capable of generating large phenotypic datasets.

We demonstrate here the application of this pipeline to CT-scans of cichlid fishes from the closely related species *Maylandia zebra* and *Cynotilapia zebroides*, which exhibit variable morphology in the jaw and head. In section 2 we first introduce this dataset and then describe the methods used in the software, with results summarised in section 3. The traits measured using the method were used to identify candidate genetic loci underlying craniofacial variations which we discuss in sections 3 and 4.

## 2 Materials and Methods

### 2.1 Malawi cichlid dataset

Lake Malawi, the southernmost great lake in the East African Rift (Ebinger, 2005), is home to one of the most diverse vertebrate radiations in the world, with more than 500 endemic cichlid fish taxa that all are believed to have diverged within the last million years (Johnson et al., 1996; Malinsky et al., 2018; Svardal et al., 2019). Despite this short diversification time, the morphological and ecological variation between species is extensive (Kornfield and Smith, 2000), and the mechanisms through which this diversity has emerged have been the subject of many studies (Coyne, 1992; Schluter, 2009; Malinsky et al., 2018; McGee et al., 2020; Nosil et al., 2021). However, while there has been increasing use of large scale sequencing data in these studies (Malinsky et al., 2018; McGee et al., 2020) so far few analyses have combined dense genetic data with detailed quantitative morphology information (Ronco et al., 2021).

Here we focus on two closely related species of rock-dwelling cichlids from the Mbuna group of species in Lake Malawi: *Maylandia zebra* and *Cynotilapia zebroides*. These are superficially similar (Figure 1 (A)), but have different adult teeth and exhibit more subtle differences in other features, in particular in their jaw and craniofacial structures. An attractive approach to obtain highly precise quantitative measurements of these features is micro-CT scanning, which can give exquisitely detailed 3D information on the underlying skeletal structure (Figure 1 (B)).

**Figure 1:**
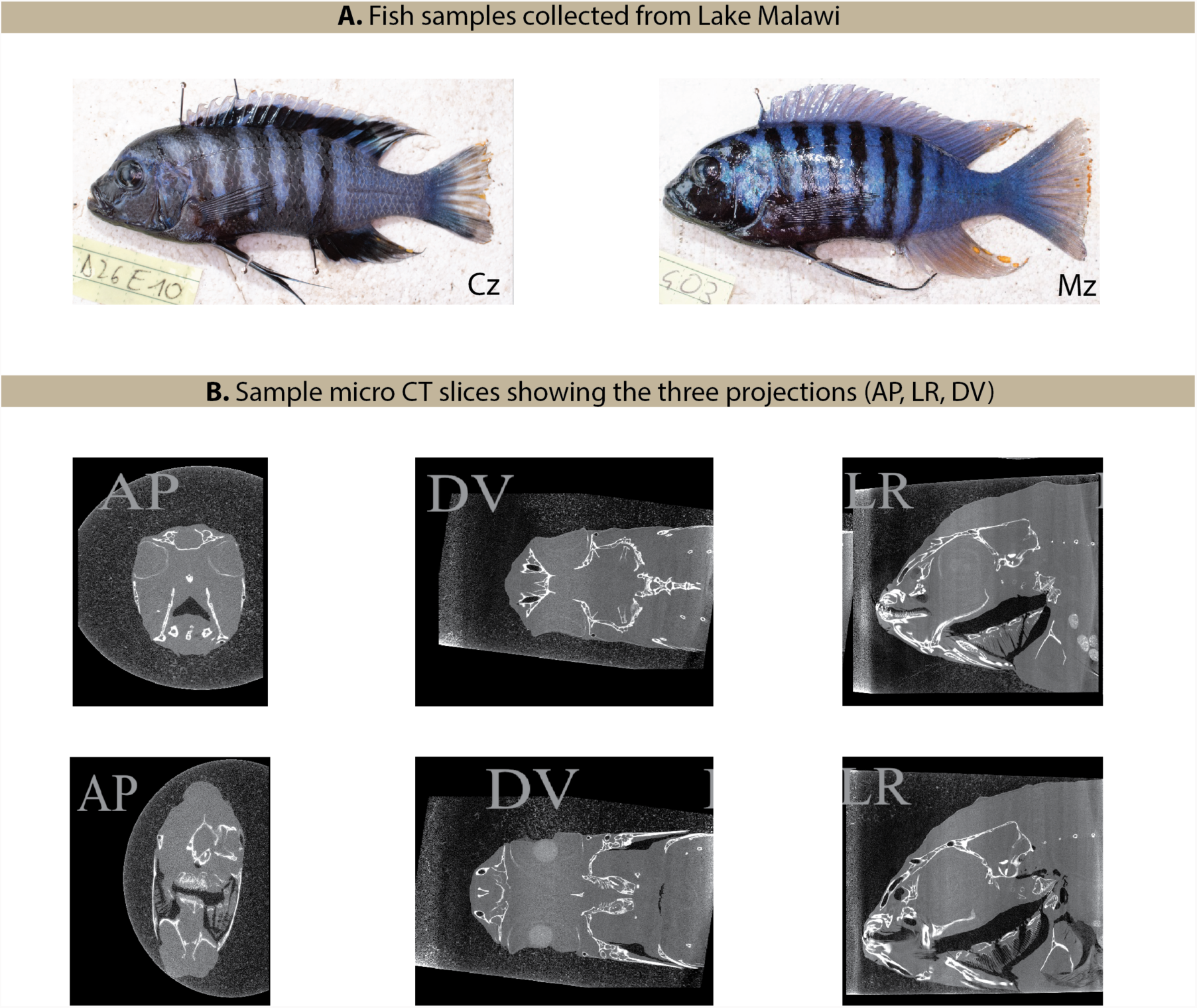
(A) Images of individual *Maylandia zebra* (MZ), and *Cynotilapia zebroides* (CZ) fish. (B) Sample CT scans of *Maylandia zebra* individuals perpendicular to the three primary axes: Anterior-Posterior (AP), Dorsal-Ventral (DV), and Left-Right (LR)

In total a set of 118 (115 after removal of 3 outliers) fish from four locations were used for this study that were collected in two field trips to Lake Malawi in 2016 and 2017 (see Table Supplementary 1 for details). These were fixed in formaldehyde and ethanol, and scanned at the Cambridge Biotomography Centre using a Nikon XT 225 ST micro-CT scanner. Tomographic reconstruction from the raw images generated stacks of 2D images with a separation of 10*µ*m. Whole genome shotgun DNA sequencing data were also collected on an Illumina HiSeq X instrument.

### 2.2 Pre-processing

The Malawi cichlid CT image dataset consists of stacks of 2D images that are re-constructed with a uniform separation specified in the metadata files generated by the scanner. These tomography maps are commonly presented in the anterior-posterior (AP) projection, however, they can be used to generate three sets of image stacks for each sample, one for each projection: Left-Right (LR), Anterior-Posterior (AP), and Dorsal-Ventral (DV) - as shown in Figure 1 (B). These images are commonly stored as 16-bit tiff files and can easily occupy tens of gigabytes per sample (mean 24*GB* per sample for our dataset). The orientation of the body, amount of blur, and noise are also all dependent on scanner settings and can vary significantly between scans taken on different days. An example of such imperfections in the scans is shown in Figure 2 (C) with the noise highlighted in red. In order to address these issues, we first implemented a pre-processing pipeline, highlighted in red in Figure 2 (A), to compress, denoise, and normalise all images in the dataset.

**Figure 2:**
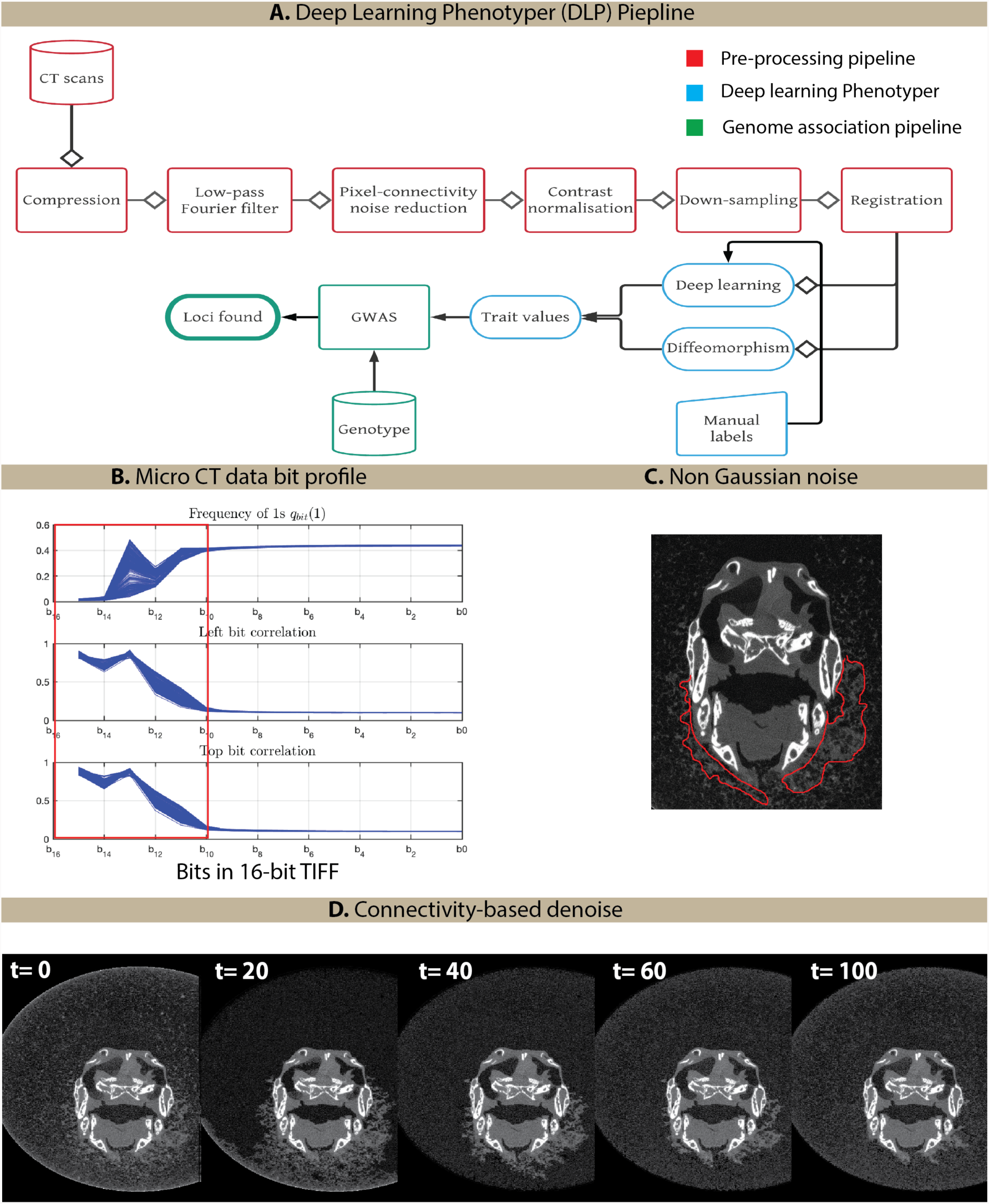
(A) Deep learning phenotyping pipeline with pre-processing steps shown in red. (B) Variation in the frequency of ”1s” in bits as well as spatial correlation plotted as a function of bit depth with the red window showing the least significant bits that can be truncated with minimal information loss. (C) Sample CT scan showing non-additive, spatially correlated aberrations in red. (D) Image stack showing how the efficacy of the pixel connectivity-based filter varies with the choice of binary thresholding value *t* ∈ {0, 20, 40, 60, 100}) from left to right.

#### 2.2.1 Compression

Image stacks are first compressed using a dynamic bit-wide compression algorithm that truncates bits showing random behaviour when looking at the ensemble of pixels in each of the images (Figure 2 A). The degree of randomness of each bit is determined using the frequency of 1s as well as the spatial correlation of the bit with the same bit of neighbouring pixels (see Section 4 for a description on how these spatial correlations were computed). Figure 2 (B) shows how the frequency of ones and spatial correlations of bits vary as a function of bit depth. The compression method implemented in the pre-processing pipeline compresses images by truncating bits (shown in red in Figure 2 B) with a compression ratio of 10-12 depending on bit profiles. Images are then stored using lossless Joint Photographic Expert Group (JPEG) format for further lossless compression (Matsuda et al., 2005).

#### 2.2.2 Noise reduction

Compressed images are then passed through a low-pass Fourier filter implemented as a fifth-order Butterworth filter (Butterworth et al., 1930) to suppress any white additive noise at frequencies higher than the image bandwidth (50 Hz), determined empirically by investigating ensembles of Fourier transforms. While white additive noise can be removed using linear Fourier filters, aberrations that are spatially correlated can not be removed, such as those highlighted in red in Figure 2 (C). A pixel-connectivity denoising filter was, therefore, implemented to remove these artefacts based on the contrasts between pixels with six-connectivity in three dimensions. Algorithm 1 describes how this filter was implemented.

##### Algorithm 1

Connectivity-based denoising

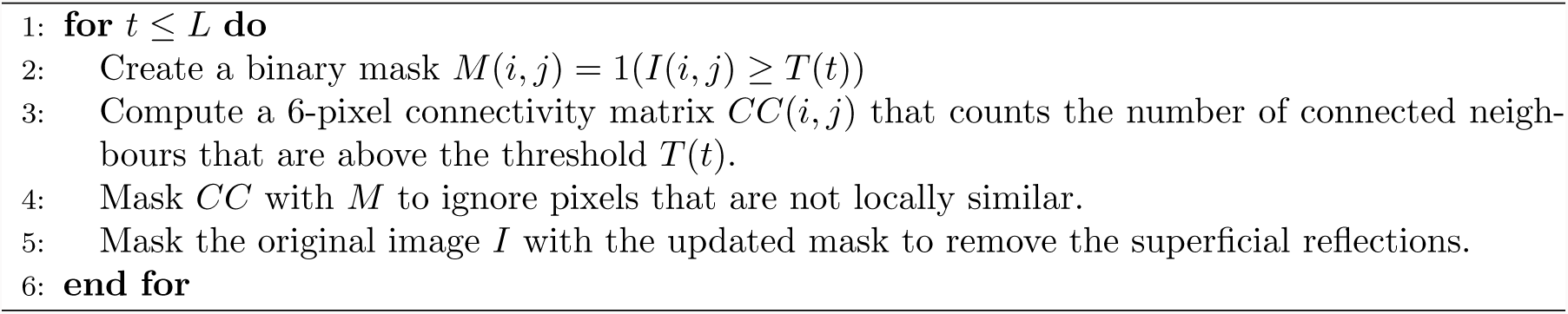

The value of *t* was iterated over a range of pre-determined values stored in an array *T* of length *L*. An optimal value of 60 was chosen by empirically investigating a series of 100 images. Figure 2 (D) shows how the efficacy of this filter in removing the aberrations varies with the choice of binary mask thresholding value.

#### 2.2.3 Contrast normalisation and down sampling

Denoised image stacks are passed through a contrast normalisation in order to normalise the contrast between images of samples scanned with different scanner settings specified in the *xteckt* files. The normalisation process is done through histogram matching, with one of the samples selected as a reference. The contrast normalisation is then followed by a Gaussian-based convolutional down-sampling step in order to normalise the dimension of all image stacks from different samples. These pre-processing methods are crucial to ensure that the variations registered using the phenotyping platform are not due to differences in scanner setup and are true measurable variations between samples. Images passed through this pre-processing pipeline were then used for phenotyping as shown in blue in Figure 2 (A).

### 2.3 Deep learning 3D phenotyper

We adopted a transfer-learning classification-based approach to phenotype morphological features from the image stacks pre-processed as discussed above. Our strategy was to train classifiers for whether or not a 2D slice of the data contained a 3D feature, such as the eye or pharyngeal jaw, and then use these to measure a bounding box for the feature with lengths along each of the three axes as in 3 (A). We call this system the Deep Learning Phenotyper (DLP).

To train the classifiers, a manually labelled training dataset of 10,000 2D images was generated for each feature and for each of the three standard dimensions, by selecting fish randomly and assigning blocks of slices that contained the feature to the set until a sufficient number of images were obtained. These training sets were then randomly shuffled and an equal number of negative images were selected at random from the remaining slices in the same fish. Labelled images were then fed into the following pre-trained networks: *Alexnet*, *Resnet50*, *Resnet18*, and *GoogleNet* (Krizhevsky et al., 2012; Szegedy et al., 2014; He et al., 2015b,a) respectively. Multiple feature spaces were generated using different pooling layers of these networks, and were then used to train a linear Support Vector Machine (SVM) (Boser et al., 1992) using the manually annotated images. Evaluation on a held out sample of labelled images identified the last average pooling layer of *Resnet50* as the optimal architecture for the task, which was consistent with previous studies showing high efficacy of *Resnet50* in CT datasets (Peng et al., 2020). The pre-trained *Resnet50* network was therefore used to generate a 2048-dimensional feature vector for every image in the image stack for individual fish scans along a given projection. Figure 3 (B) shows the first two Principal Components (PCs) of the training feature space for a binary classifier of an eye/ no-eye classification.

**Figure 3:**
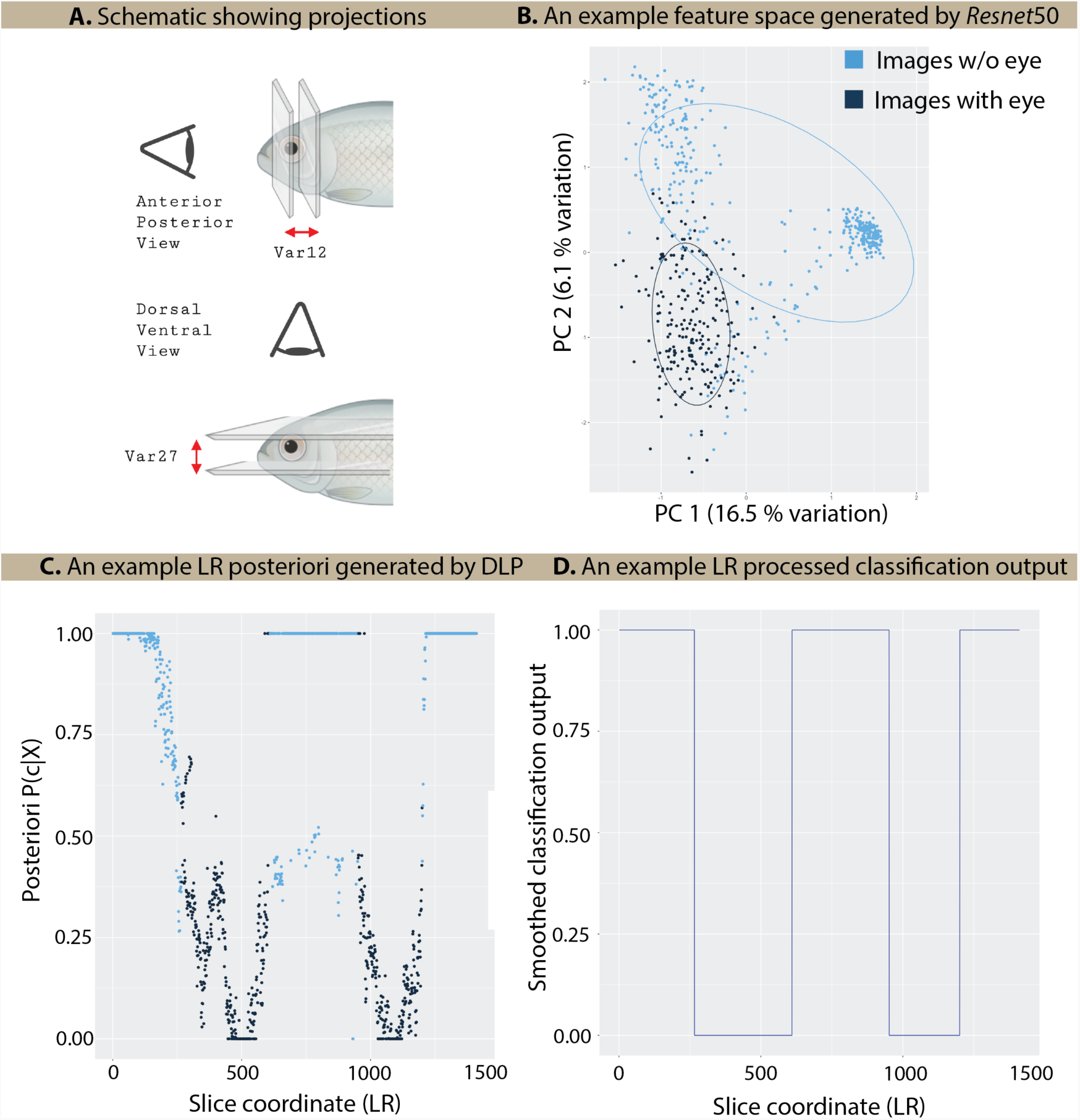
(A) Schematic diagram showing how classification of major anatomical feature (e.g the eye) can be used to determine how wide, deep and high that anatomical feature is using all three projections (AP, LR, and DV). (B) The first two Principal Components (PCs) of the LR feature space generated by the last average pooling layer of *Resnet50*. The colours correspond to different labels on the images (i.e eye or no eye classes) - (C) Posteriori *P* (*c|X*), generated by the DLP for images in the LR projection of a sample fish, plotted against the LR axis coordinate of the body. (D) Label vector filtered for abnormalities in classification using the smoothing algorithm given in Algorithm 2

We next evaluated a range of SVM kernel functions for classification over this feature space, using 60% of the annotated dataset for training (7,500 images) and 20% for model selection (2,500 images). Using this approach, a linear SVM model was shown to outperform all the other models for all three projections (AP, LR, and DV). The confusion matrices and classification accuracy given in the Results section were then calculated using the final 20% of the evaluation data which had not been previously used in the training and optimisation process. Images that are not labelled are first passed through *Resnet50* to encode them each into a 2048-dimensional vector. These vectors are then passed through the trained SVM models to classify each image, with the output providing a posterior probability of the slice containing the major craniofacial feature that the SVM was trained on. These values are then plotted along each of the three projections in order to create “projection profiles” that indicate how long, wide, and deep each of the labelled anatomical features is (Figure 3 C).

Due to noise and other adverse effects, the SVM classifier can have uncertain posterior predictions and even misclassify some of the images (Figure 3 C). To address this, the posterior probabilities in a profile are passed through a smoothing algorithm that leverages the continuity of anatomical features to correct misclassified images, as shown in Algorithm 2. This algorithm also returns the left- and right-most positions at which the posterior probability is above the 60% threshold. These boundaries are used to calculate a potential error measure for the feature. Figure 3 (D) shows the output of this algorithm applied to the initial SVM output shown in Figure 3 (C).

#### Algorithm 2

Classification error removal

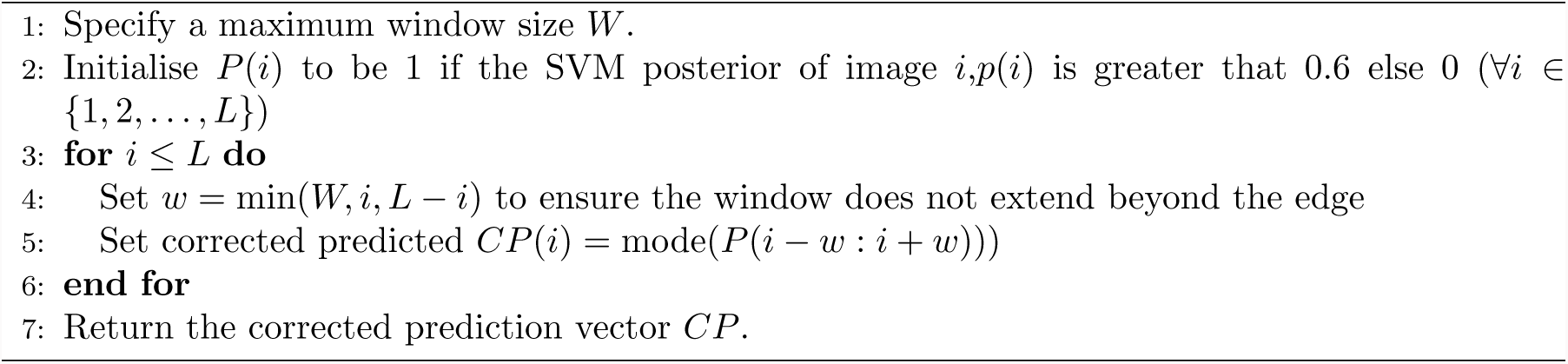

The final step in the pipeline (Figure 2 A) is a conversion step that maps the number of consecutive images labelled to contain the same anatomical feature to the physical length of that feature along the current axis by multiplying by the voxel size along that axis.

### 2.4 Genetic analysis

We identified candidate genetic loci associated with variation in the measured traits of 61 *M. zebroides*and 54 *C. zebroides* (after removing three outliers as described below) using the generalized linear modeling framework of ANGSD (Korneliussen et al., 2014). This approach accounts for genotyping uncertainty through incorporation of genotype posterior probabilities in the association likelihood model. Following standard quantitative genetics approaches, ANGSD models the traits **y** = (**y**_1_, …, **y**_115_) of the 115 individuals as normally distributed, such that for the *i*’th individual 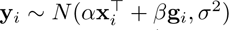. The genotype **g***_i_* ∈ {0, 1, 2} denotes the number of minor alleles at a given SNP and **x***_i_* = (1, **x***_i_*_1_, …, **x***_i_*_8_) is a vector of covariates that control for variables that may confound causal genetic associations. We used the length of the head (**x***_i_*_1_) to account for the overall size of the fish and scores (**x***_i_*_2_, …, **x***_i_*_8_) of the first seven principal components (PCs) of a principal component analysis (PCA) of genome-wide genetic variation to control for population structure. We fit global maximum likelihood estimates of 𝛼 = (𝛼_0_, 𝛼_1_, …, 𝛼_8_), and then for each locus compared the maximum likelihood with 𝛽 free, allowing for uncertainty in the genotypes using the genotype posterior probabilities, to the likelihood when 𝛽 = 0, which corresponds to the null hypothesis of no genetic effect, using a likelihood ratio test (Skotte et al., 2012).

We tested for genetic trait associations at 2,693,904 biallelic SNPs having a minor allele frequency (MAF) of at least 2% and for which at least 10% of individuals had a posterior probability of 0.9 or greater for at least two of the three possible genotypic configurations (0, 1, or 2 minor alleles). We also required that at least five individuals had a called genotype with quality of at least 15 using GATK (McKenna et al., 2010), and that at least two of the three genotypic configurations were represented in this manner at each tested SNP. We calculated a p-value of genetic association for each SNP by comparing twice the log likelihood ratio to a 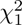-distribution, which we considered significant if it exceeded 5% after a conservative Bonferroni correction for the number of tested SNPs (−log10(p-value) *>* 7.73).

The PCA of genome-wide genetic variation used to control for population structure in the GWAS was performed using ngsTools (Fumagalli et al., 2014) and R. Genotype likelihoods for all individuals calculated with ANGSD were used with ngsTools to estimate a genetic covariance matrix from SNPs with a likelihood ratio test p-value for being variable below 1e-6 and minimum MAF of 5%. This covariance matrix was decomposed using the ’eigen’ function of R, yielding the eigenvectors used to control for population structure. We used the first seven PCs as covariates in the GWAS since these captured the genetic structure between the species and sub-populations within species (see Figure 5 B for the first two PCs).

### 2.5 Implementation and availability

The main package for DLP and pre-processing is written in MATLAB 2021R, with ancillary scripts in R and bash. All of the code required to implement DLP and also the R scripts used to generate the plots in Figures 3, 4, and 5 are available along with documentation at **(removed for anonymous peer review)**. This documentation guides users through pre-processing images before labelling and training the SVM. There is also guidance and scripts for users to integrate the package with ANGSD if they seek to run genetic association tests.

## 3 Results

### 3.1 Phenotyping method

Our goal was to implement an automated package that would require minimal manual input to ease the phenotyping process for making skeletal measurements from micro-CT data, while keeping a high level of accuracy. To this end, we first explored three diffeomorphic mapping algorithms (Ashburner and Friston, 2000; Friston et al., 2007; Ashburner, 2007) to register variations in morphology to a reference image (Figure 2 (A)). However, none of these were successful, we believe due to not having an adequate reference atlas as well as noise and aberrations in the scans. The variational methods we investigated had previously been applied to Magnetic Resonance Imaging (MRI) scans (Friston et al., 2007), for which high-resolution atlas datasets exist and all input images are extensively manually pre-processed.

We next considered deep learning methods, due to their ability to generalise well when trained with care to avoid over-fitting. Recent developments in image classification and segmentation using deep learning have resulted in comparable performance to manual classification and segmentation (LeCun et al., 2015; Mukhoti and Gal, 2018; Grigorescu et al., 2019). Within deep learning methods, we first attempted to extract phenotypic measurements of the skull by segmenting the CT images, however, the performance of segmentation implemented using a variety of deep convolutional neural networks (Havaei et al., 2015; He et al., 2015a; Chen et al., 2018) was poor. We then investigated architectures developed for object detection(Fernández et al., 2007; Redmon et al., 2015; Redmon and Farhadi, 2016, 2018) to localise biologically significant bones, fragments or other anatomical features within the skull. While the performance improved over segmentation approaches with the same level of manual training information, the overall measurement accuracy was still not acceptable, with under 50% overlap between the predicted bounding boxes and the manually annotated ones in the evaluation dataset.

To resolve these issues, we decided to formulate the phenotyping problem as a classification task. More specifically, we used a classification network to assess whether or not a given anatomical feature was present in a given two dimensional slice of the data. By applying this to all the slices along an axis, for example the Anterior-Posterior (AP) axis, we could identify intervals along that axis that contain the feature, and hence measure its length along that axis. By carrying out this procedure on image stacks generated for each of the AP, Left-Right (LR) and Dorsal-Ventral (DV) axes we could measure the size of features such as bones in each of the three canonical orthogonal directions. Figure 3 (A) shows an example of how classification can be used in two different projections to measure the eye diameter of a fish.

This classification approach allowed us to use transductive transfer learning approaches (Pan and Yang, 2010; Weiss et al., 2016) that are commonly implemented using very well-established networks (Krizhevsky et al., 2012; Szegedy et al., 2014; He et al., 2015b), trained on very large annotated image datasets (see (Lin et al., 2014; Russakovsky et al., 2015) for examples of these datasets). Many different network architectures have been proposed in the last few years with astonishingly high accuracies in image classification, some beyond human performance. It is common lore that the first layers of deep CNNs trained on large datasets extract morphological features that are essentially independent of the image dataset used. This suggests that a much less complex model could be trained if only the final few layers of these deep networks were to be trained on the new dataset, as opposed to training millions of parameters from scratch.

We implemented this approach as described in detail in the Methods section 2.3. There are two steps in how we extract phenotypic data from images using CNN classifiers. The first step trains a support vector machine (SVM) on the features generated by the last pooling layer of the *ResNet50* network (He et al., 2015b). Figure 4 panels (A, B, C) show confusion matrices for classification of slices by SVMs for each of the AP, LR and DV axes, evaluated on held out labelled data using a 50% posterior probability threshold. These give overall SVM classification accuracies of 94%, 92% and 89% for the three axes respectively. A second step smooths these posterior estimates across slices to give a continuous interval along the axis that is predicted to contain a given feature (as in Figure 3 D), then converts the length of this interval into a physical length using the voxel spacing. Training was fast, taking approximately ten minutes on a single cluster node with four GPU cores.

**Figure 4:**
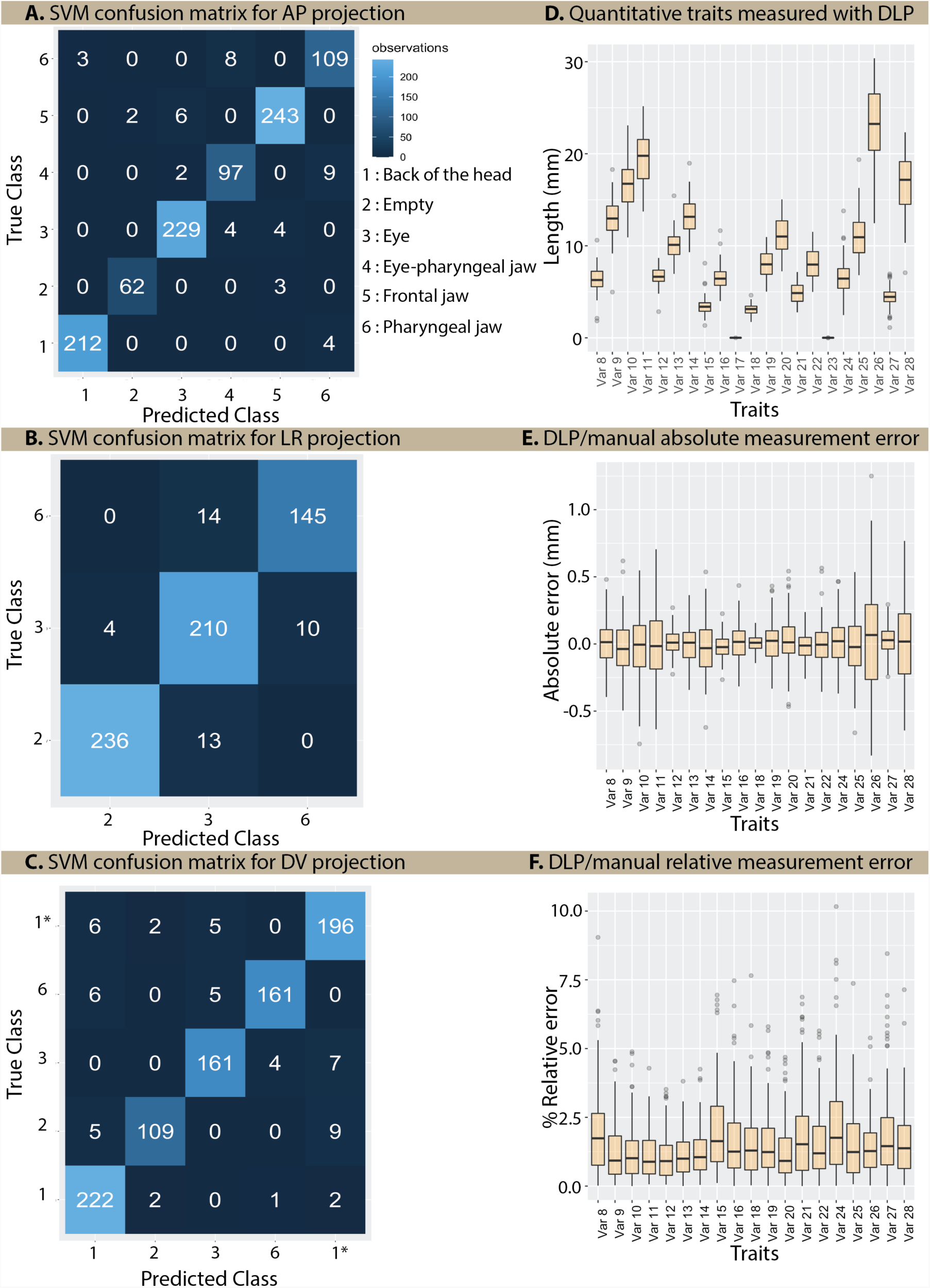
(A, B, C) confusion matrices for classification of slices by SVMs for each of the AP, LR and DV axes, evaluated on held out labelled data using a 50% posterior probability threshold. These give overall SVM classification accuracies of 94%, 92% and 89% for the three axes respectively. (D) Boxplot showing distributions for 19 traits and 2 controls that were measured using the DLP. The error in measurement was assessed by comparing the DLP measurements with measurements that were conducted manually using the Slicer interactive software (Kikinis et al., 2014). (E) and (F) show these error values both in absolute mm and relative to the size of the respective measurement.

### 3.2 Craniofacial trait measurement

The pipeline described above was applied to the dataset of 61 *M. zebroides* and 54 *C. zebroides* micro-CT scans (see Methods section 2.1). Twenty-one different measurements were made as listed in Table 1. Nineteen of these are quantitative craniofacial traits, while two measures, Var17 and Var23, correspond to the maxima of the error measures across all features for the LR and DV projections.

**Table 1:** Description of the traits extracted by DLP with their summary statistics.

| Trait | Description / Projection | Mean (mm) | Standard deviation (mm) |
| --- | --- | --- | --- |
| Var8 | Length of the frontal jaw / AP | 6 | 1.34 |
| Var9 | Length of the Preorbital bone / AP | 13.0 | 2.13 |
| Var10 | Length of the Posttympanic bones / AP | 16.4 | 2.46 |
| Var11 | Total length of the head / AP | 19.4 | 2.91 |
| Var12 | Eye diameter / AP | 6.69 | 1.39 |
| Var13 | Length of the Posttympanic bone / AP | 10.1 | 1.45 |
| Var14 | Eye to back of head separation / AP | 13.1 | 1.92 |
| Var15 | Length of the pharyngeal jaw / AP | 3.44 | 0.87 |
| Var16 | Pharyngeal jaw to back of head separation / AP | 6.45 | 1.23 |
| Var17 | Error (resolution) in the LR projection | 4.18E-5 | 1.45E-5 |
| Var18 | Depth of the first eye / LR | 3.11 | 0.57 |
| Var19 | Width of the Preorbital bone - left / LR | 7.98 | 1.46 |
| Var20 | Total width of the head / LR | 11.1 | 1.99 |
| Var21 | Length of the pharyngeal jaw / LR | 4.87 | 1.07 |
| Var22 | Width of the Preorbital bone - right / LR | 8.00 | 1.58 |
| Var23 | Error (resolution) in the DV projection | 4.36E-5 | 1.45E-5 |
| Var24 | Height of the Frontal bone / DV | 6.60 | 1.70 |
| Var25 | Height of the Preorbital bone / DV | 11.0 | 2.15 |
| Var26 | Total height of the head / DV | 23.3 | 3.96 |
| Var27 | Depth of the eye / DV | 4.44 | 0.99 |
| Var28 | Height of the Preopercle bone / DV | 16.7 | 3.10 |

Figure 4 (D) shows the size range of measurements for each feature. The error in measurement was assessed by comparing the DLP measurements with measurements that were conducted manually using the Slicer interactive software (Kikinis et al., 2014). Figure 4 panels (E) and (F) show these error values both in absolute mm and relative to the size of the respective measurement. Absolute errors are typically below 0.1mm, mostly within 1% of the trait value. This is substantially below the standard deviation of trait values seen across individuals, as seen in Figure 4 (D) and Table 1, indicating that the automated DLP method generates informative measurements for downstream analysis of inter-individual variation. We note that the error values presented here, which are several times the slice resolution of 10*µ*m are upper estimates, because there is likely to be error in the manual measurements using Slicer.

### 3.3 Illustrative use for quantitative genetics

To illustrate the utility of the DLP measurements for downstream analysis, we present here an initial GWAS analysis of these samples using general linear models. Figure 5 (A) shows the first two principal components (PCs) of the nineteen traits, colour coded by species. It is apparent that PC1 largely separates the samples by species. PC2 separates out three samples from the rest, which we treat as outliers and removed from the association tests. We note that in general the *M. zebroides* samples are larger than the *C. zebroides* samples, so that all measurements tend to be larger, although the relative differences vary between traits (Figure Supplementary 1) and indeed PC1 is correlated to overall size as measured by the head length (Var11) with correlation coefficient of 0.49.

**Figure 5:**
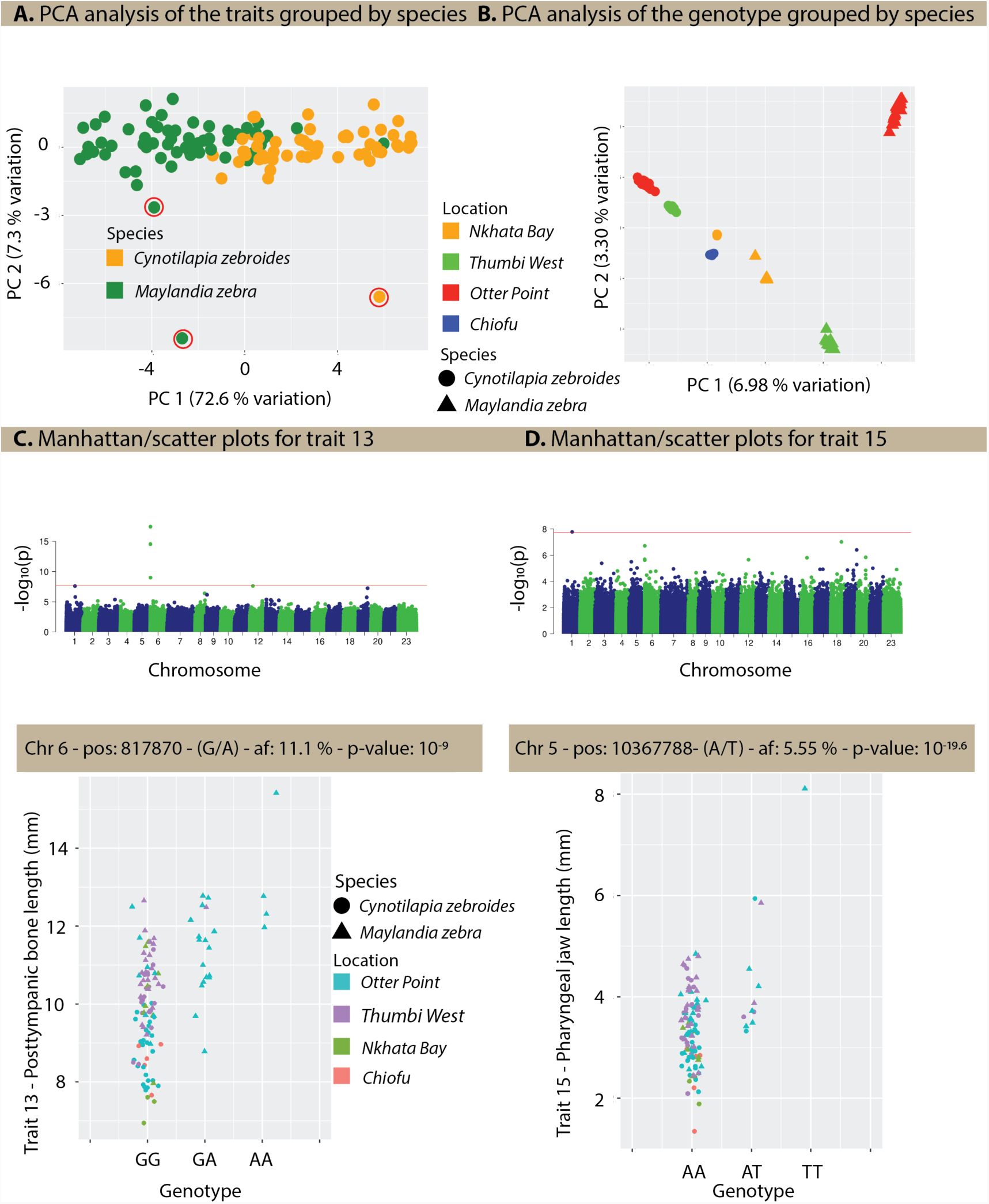
(A) The first two principal components of the quantitative traits measured by the DLP with points labelled according to species. The individuals shown in red are the outliers that were excluded from the GWAS. (B) The first two principal components from a PCA of genome-wide SNP variation (see section 2.1). (C) Manhattan and phenotype-genotype scatter plots showing correlations for traits 13 (Posttympanic bone length) for the genome-wide scans of association between SNPs and trait 13. In the Manhattan plots each point indicates the p-value of a SNP, with alternating blue and green colours denoting SNPs on different chromosomes. The red, horizontal line shows the 5% genome-wide significance level after a Bonferron1i 5adjustment for multiple testing. (D) Manhattan and phenotype-genotype scatter plots showing correlations for traits 15 (Posttympanic bone length)

The primary axis from a PCA of genome-wide variation using 926,575 SNPs with a minimum MAF of 5% describes genetic differences between species, while PC2 explains differences mainly between Thumbi West and the Otter Point populations of *M. zebroides*. Individuals of both species form distinct clusters based on locality along PCs 1 and 2, indicating clear genetic population structure according to species and geography. We controlled for spurious genetic associations arising from background genetic population structure by including the genetic PCs (1-7) that capture this population substructure as covariates in the GWAS analysis, also including Var11 (head length) as a covariate to control the systematic differences described in the previous paragraph.

As a control, we first performed a GWAS on the two technical measurement error estimates from the morphometric analysis (Var17 and Var23), for which there should be no genetic association. Indeed, neither of these genomic scans identified any significant loci. Association scans for seven of the remaining 18 traits (Var11 was used as a covariate and not tested) yielded nominally significant genetic associations across a total of 33 loci (Table 2, see Figure 5 for examples of significant associations with Posttympanic bone length (panel C) and pharyngeal jaw length (panel D)). There is substantial overlap in significant SNPs across different traits, with 5 : 10367788 associated with six traits. This is perhaps not surprising since the traits are not independent. Inclusion of the genetic PCs and head-length covariate effectively controlled for p-value inflation (Figures Supplementary 2 - Supplementary 6), offering support for the authenticity of the identified associations.

**Table 2:** Statistically significant loci found for traits 8,9,12,13,14,15 and 16 with the highlighted SNPs corresponding to the loci pictured in the phenotype-genotype plots of Figure 5 (C) and (D)

| Chr | Position | Frequency | Major | Minor | Traits | $-\log_{10} p$ values |
| --- | --- | --- | --- | --- | --- | --- |
| 1 | 28508896 | 9.9% | C | A | 8,14 | (157,154) |
| 1 | 28508919 | 10.0% | T | G | 15 | 27 |
| 2 | 12359064 | 27.6% | A | G | 9,16 | (8,8) |
| 3 | 7173111 | 5.5% | C | T | 12 | 45 |
| 3 | 9058204 | 11.7% | A | T | 8,14 | (12,12) |
| 3 | 14695098 | 6.5% | A | G | 9,16 | (31,31) |
| 3 | 48468960 | 5.5% | C | T | 9,16 | (75,76) |
| 5 | 10367788 | 5.5% | A | T | 8,9,13,14,15,16 | (17,13,13,16,20,13) |
| 5 | 35205044 | 44.5% | T | A | 12 | 13 |
| 6 | 709539 | 10.2% | C | T | 8,13,14 | (7,17,7) |
| 6 | 817870 | 11.1% | G | A | 13 | 9 |
| 6 | 5698280 | 5.4% | C | T | 9,16 | (98,100) |
| 7 | 8753577 | 7.8% | C | T | 12 | 11 |
| 7 | 18398467 | 5.9% | T | C | 8,14 | (31,31) |
| 7 | 25825327 | 7.2% | C | T | 8,13,14,15 | (86,200,85,9) |
| 7 | 67050617 | 6.7% | G | A | 12 | 24 |
| 9 | 22036037 | 5.9% | G | A | 8,13,14,15 | (63,229,62,19) |
| 10 | 33712863 | 5.3% | A | G | 12 | 40 |
| 11 | 29806372 | 5.4% | T | C | 9,16 | (273,284) |
| 13 | 5331913 | 9.6% | T | A | 8,13,14 | (9,25,9) |
| 14 | 8941641 | 10.7% | G | A | 9,16 | (9,9) |
| 14 | 15181937 | 7.3% | T | C | 8,13,14,15 | (27,33,27,12) |
| 15 | 25934106 | 23.0% | A | G | 8,14 | (11,11) |
| 15 | 27258227 | 5.6% | A | G | 9,16 | (31,31) |
| 15 | 29560314 | 5.5% | G | A | 9,16 | (210,217) |
| 17 | 7685438 | 5.4% | C | A | 12 | 15 |
| 19 | 5701806 | 5.9% | C | A | 9,16 | (48,48) |
| 20 | 21728641 | 10.7% | A | G | 9,16 | (8,8) |
| 22 | 2316979 | 5.4% | G | A | 9,16 | (27,27) |
| 23 | 103914 | 5.3% | C | A | 8,13,14 | (17,11,17) |
| 23 | 16225831 | 9.8% | T | C | 15 | 78 |
| 23 | 31412637 | 5.5% | G | A | 15 | 76 |

## 4 Discussion

The DLP package was developed to enable fast and accurate morphometric phenotyping that is almost entirety automated, and therefore is capable of scaling to large sample sizes. We have shown that it is accurate when compared to time-consuming and intrinsically more subjective manual approaches. It is also robust in that only three out of 118 samples had to be rejected as outliers. It requires relatively small amounts of training data and is fast to train and apply to new samples.

Performing morphological analysis through the classification of slices in micro-CT data is significantly easier and more accessible to implement compared to existing land-marking based techniques in three dimensions (Webster and Sheets, 2010; Gunz and Mitteroecker, 2013; Burress, 2016; Pauers et al., 2018; Hendrick et al., 2019) since the slice-based labelling process for training only takes about an hour to perform for all three projections for each 3D feature. The models trained using these labelled images can then be applied to an entire micro-CT dataset, and images from each sample do not need to be manually processed or land-marked before phenotyping.

To showcase how these measurements can be used, we trained the DLP network to investigate the morphological variation that is observed both within and between two closely related cichlid fish species: *Maylandia zebra* and *Cynotilapia zebroides*. These measurements were then used in a GWAS for 18 of the traits in order to identify candidate genetic loci involved in determining craniofacial morphology. No significant results were found using two sets of control traits, whereas for seven skeletal traits we found statistically significant genetic associations at 33 loci. These loci are promising, but particularly in light of the small sample size for GWAS should be regarded as candidate loci for further investigation by replication with additional samples and functional exploration, for example through genetic editing or gene expression experiments, rather than proven biological associations.

We have shown that morphological measurement of intricate skeletal elements can be carried using deep convolutional neural networks, and that overfitting can be avoided if transfer learning approaches are used, specifically based on the powerful 2D image classification network *Resnet50*. Beyond genetic studies, the DLP package can potentially be used for other quantitative skeletal analysis applications using micro-CT data. Although we only demonstrate its use here on the heads of fish, there was no specific knowledge of fish built into the package, and so we infer that it would be applicable to other vertebrate skeletal systems, or, more generally, to other 3D datasets comprised of 2D image stacks in which 3D elements can be observed.

## Supporting information

Supplemental section and figures

## Acknowledgement

For assistance in sample collection we would like to acknowledge the Government of Malawi, additional members of the Cambridge Malawi cichlid field expeditions in 2016 and 2017: Milan Malinsky, Alix Tyer, George Turner, Mingliu Du, Mexford Mpulungu, Hannes Svardal, Gregoire Vernaz, Karl Svardal, and Emília Santos; and the local guides and fishermen who assisted in the collection. CT-scanning was performed by Keturah Smithson at the Cambridge Biotomography Centre, and DNA sequencing was carried out by the Scientific Operations core at the Wellcome Sanger Institute. Initial sequence data processing was performed by Hubert Denise. We would like to acknowledge the comments and feedback of Luka Moritz Blumer and Regev Schweiger on the genetics association study. This work was funded through grants from the Wellcome Trust (WT206194 to the Sanger Institute, WT207492 to R.D.) and a studentship from Trinity Hall (to A.K.). For the purpose of open access, the author has applied a CC BY public copyright licence to any Author Accepted Manuscript version arising from this submission.

## Conflict of Interest Statement

The authors declare no conflict of interests.

## Author Contributions

RD conceived the research problem and contributed to the method design and analysis; RZ and BR collected samples; AK developed and implemented the compression, pre-processing, and the phenotyping pipeline; TL and AK performed the GWAS; AK processed the results and led the writing of this manuscript. All authors contributed critically to the drafts and gave final approval for publication.

## Inclusion statement

This study arises from a long term collaboration between scientists in Malawi and the United Kingdom. The specific methods described in this paper were developed during a masters’ project in the UK for application to data from samples collected in Malawi. Ongoing research under an Access and Benefit Sharing agreement will involve local research students and scientists in experimental design and analysis.

## Code and Data Availability

The code can be found on the following GitHub page:

https://github.com/armankarshenas/DLP

We have deposited an example dataset in dryad with doi: https://doi.org/10.6078/D1M13B containing both the primary CT data for a sample (as an Anterior-Posterior image stack), and the denoised and compressed image stacks in all three orientations (AP, Left-Right, Dorsal-Ventral) which are generated by the initial pre-processing steps described in the paper.

## Notes

https://datadryad.org/stash/share/24DX_buyJZ4FYnhkK_pyXetNweeVpgCD46aYrQuZP6A

### Competing Interest Statement

The authors have declared no competing interest.

### Summary of Updates

Three typos were corrected in this version

