## Supplemental section and figures for "A deep learning phenotyping method for genetic analysis of 3D micro-CT data"

#### Dataset field trips and locations

The table below summarises the field trips and the locations from which the fish samples were collected.

Table Supplementary 1: Summary of the location, field trips and number of samples of different dataset blocks that were used.

| Block identifier | Location | Collection trip | Collection year | Number of samples |
|---|---|---|---|---|
| D01 | Nkhata Bay | Trip 1 | 2016 | 8 |
| D03 | Nkhata Bay | Trip 1 | 2016 | 3 |
| D09 | Chiofu | Trip 1 | 2016 | 6 |
| D23 | Thumbi West | Trip 2 | 2017 | 1 |
| D25 | Otter Point | Trip 2 | 2017 | 35 |
| D26 | Otter Point | Trip 2 | 2017 | 2 |
| D26 | Thumbi West | Trip 2 | 2017 | 24 |
| D27 | Thumbi West | Trip 2 | 2017 | 18 |
| D27 | Otter Point | Trip 2 | 2017 | 13 |
| D28 | Otter Point | Trip 2 | 2017 | 5 |
| Total | | | | 115 |

#### Spatial correlation for compression

The raw CT data was encoded in 16-bit TIFF (Tagged Image File Format) files covering an intensity range of 0 - 65,535. While a broader range for the pixel-intensity value normally corresponds to sharper images with better contrast (which is of great importance in medical imaging), it may be the case that many of the low-order bits behave randomly and do not code any real information. We proposed that the least significant bits in a 16-bit pixel intensity value behave almost randomly, and they could, therefore, be removed with no adverse effect on the contrast.

To test the hypothesis, we looked at the following measures:

- Frequency of ones in each of the bits of each pixel of each image
- Spatial correlation of the bits with the pixel to their left
- Spatial correlation of the bits with the pixel above them
- The histogram of pixel intensity across all images

The spatial correlation of the bits based on their connectivity for two pixels was computed using:

$$\rho = \frac{\sum_{i=1}^{m}(x_1(i) - \bar{x_1})(x_2(i) - \bar{x_2})}{\sqrt{\sum_{i=1}^{m}(x_1 - \bar{x_1})^2}\sqrt{\sum_{i=1}^{m}(x_2 - \bar{x_2})^2)}}, \qquad (1)$$

where $x_1$ and $x_2$ are binary vectors for all the pixels in the image with one vector being a shifted version of the other. Different spatial correlation patterns could be investigated using alternative concatenation and shifting methods.

Figure 2 (B) show an ensemble profile of bit randomness in a sample of all 230,000 images. The frequency of each bit increases up to an asymptote of approximately 0.42. A frequency of ones close to 0.5 indicates that the bits appear to take a high value at random. The asymptote is actually around 0.42, less than 0.5, because around 16% of the pixels had been set to zero.

In contrast, the spatial correlation of the bits decreases as we consider less significant bits, asymptotically approaching 0. We expect that noise will be uncorrelated, while meaningful bits will show some spatial correlation. We can see, from figure 2 (B), that almost every single image seems to have 10 redundant bits that carry close to zero contrast information. Using a threshold of 0.4 for the frequency of ones and 0.1 for the spatial correlation, bits with lower values were removed and replaced with a zero. The images were then encoded as 8-bit loss-less JPEGs (Joint Photographic Expert Group) using Huffman coding (Huffman, 1952). This resulted in a dynamic compression algorithm capable of compressing images with a compression ratio as high as 12.

### Supplementary figures

#### Traits measured by DLP

Boxplots in Figure 4 panels (D,E,F) show the traits measured by DLP, as well as the absolute and relative error when compared to manual measurement. Here, we present the same data shown in panel (D) of Figure 4, with the measurements colour-coded by their species.

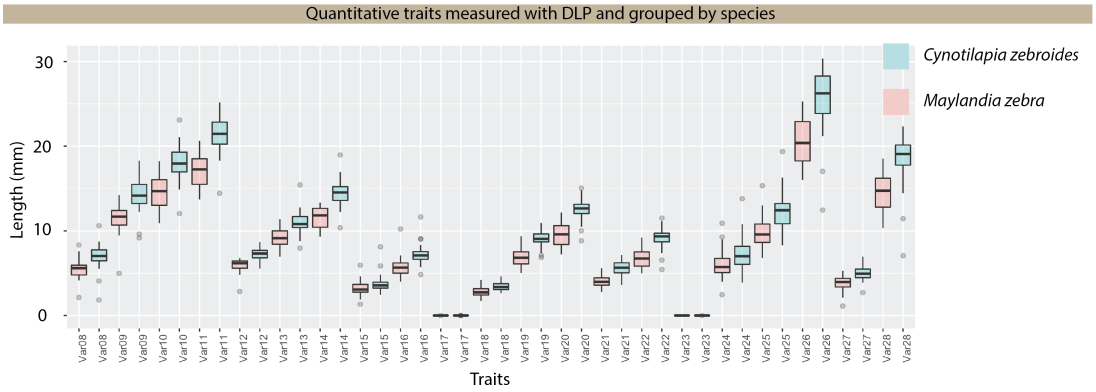


Figure Supplementary 1: Quantitative morphometric measurements done by DLP, colour-coded by species: *Maylandia zebroides* and *Cynotilapia zebroides*

#### GWAS Manhattan and QQ plots

Figure 5 (C,D) show two examples of the Manhattan and genotype-phenotype scatter plots. Here we present the Manhattan plots and QQ plots for all the traits and controls.

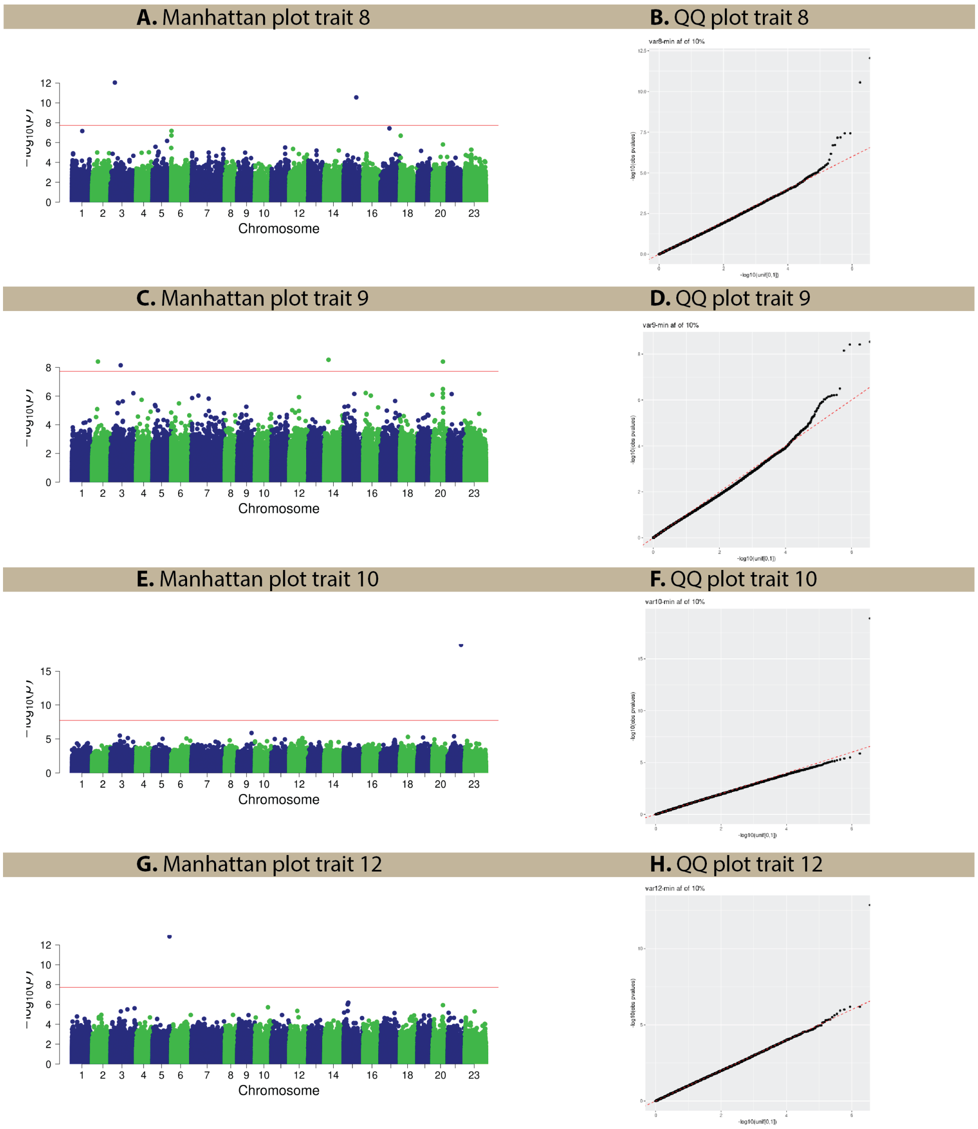


Figure Supplementary 2: (A,C,E,G) Manhattan plots for traits 8,9,10, and 12. (B,D,F,H) QQ plots for traits 8,9,10, and 12.

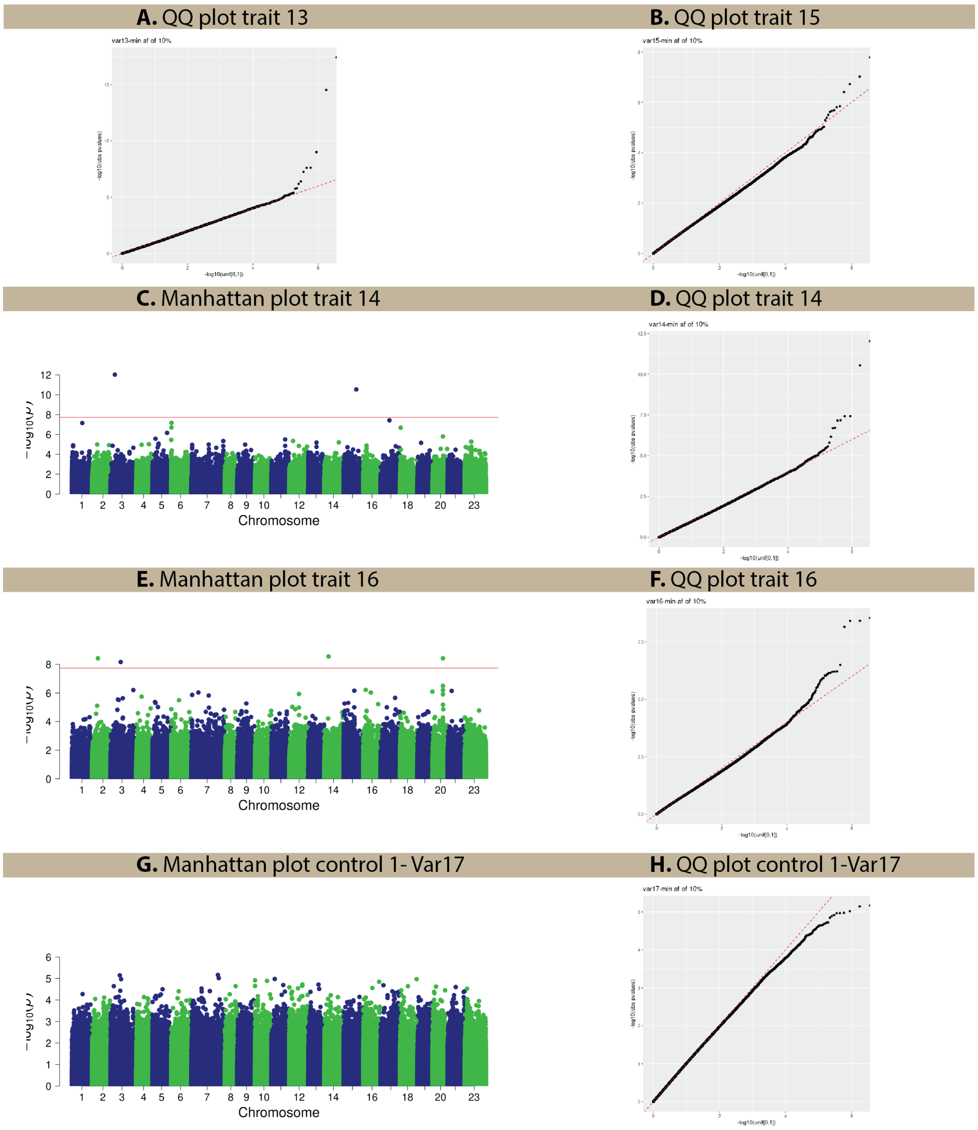


Figure Supplementary 3: (A,B) QQ plots for tratis 13, and 15 shown in Figure 5 (C,D). (C,E,G) Manhattan plots for traits 14,16, and 17. (D,F,H) QQ plots for traits 14,16, and 17

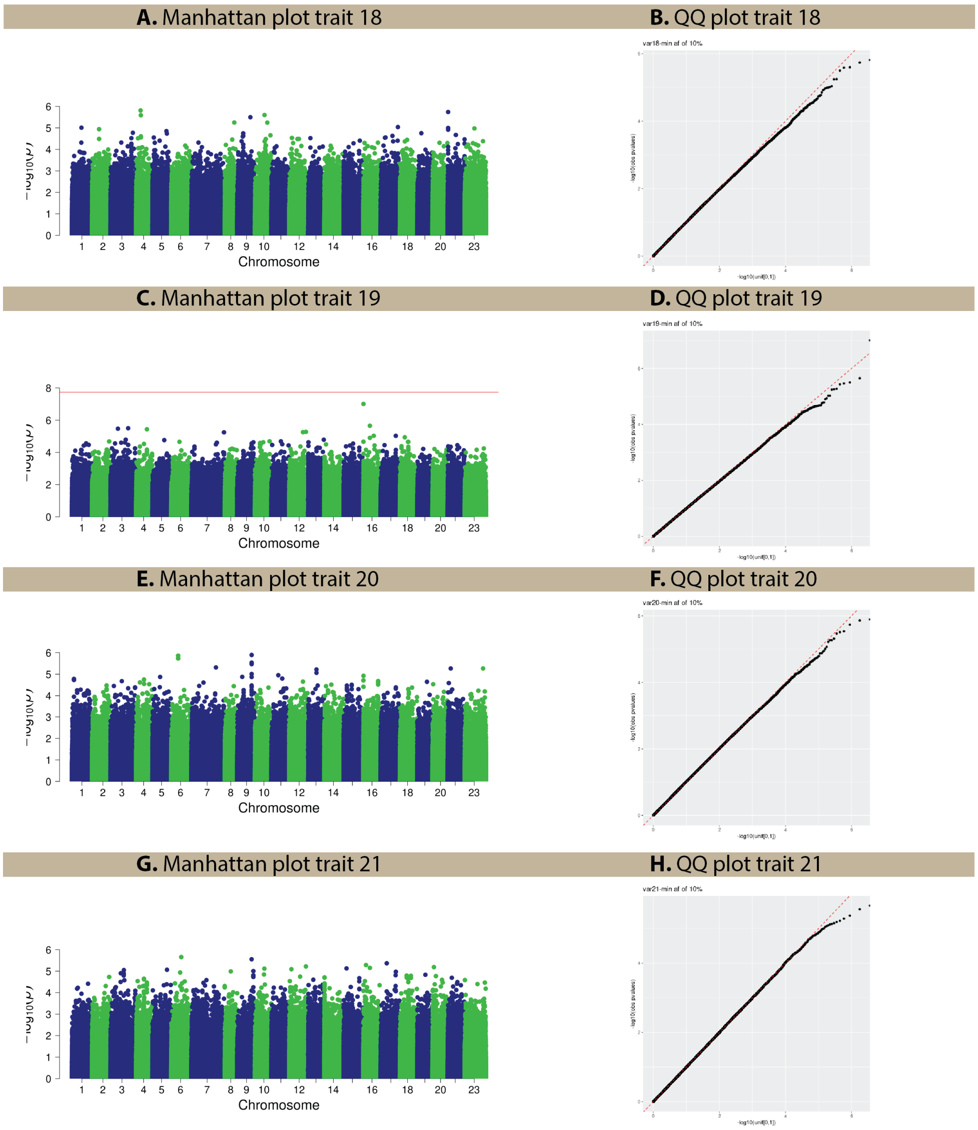


Figure Supplementary 4: (A,C,E,G) Manhattan plots for traits 18,19,20, and 21. (B,D,F,H) QQ plots for traits 18,19,20, and 21.

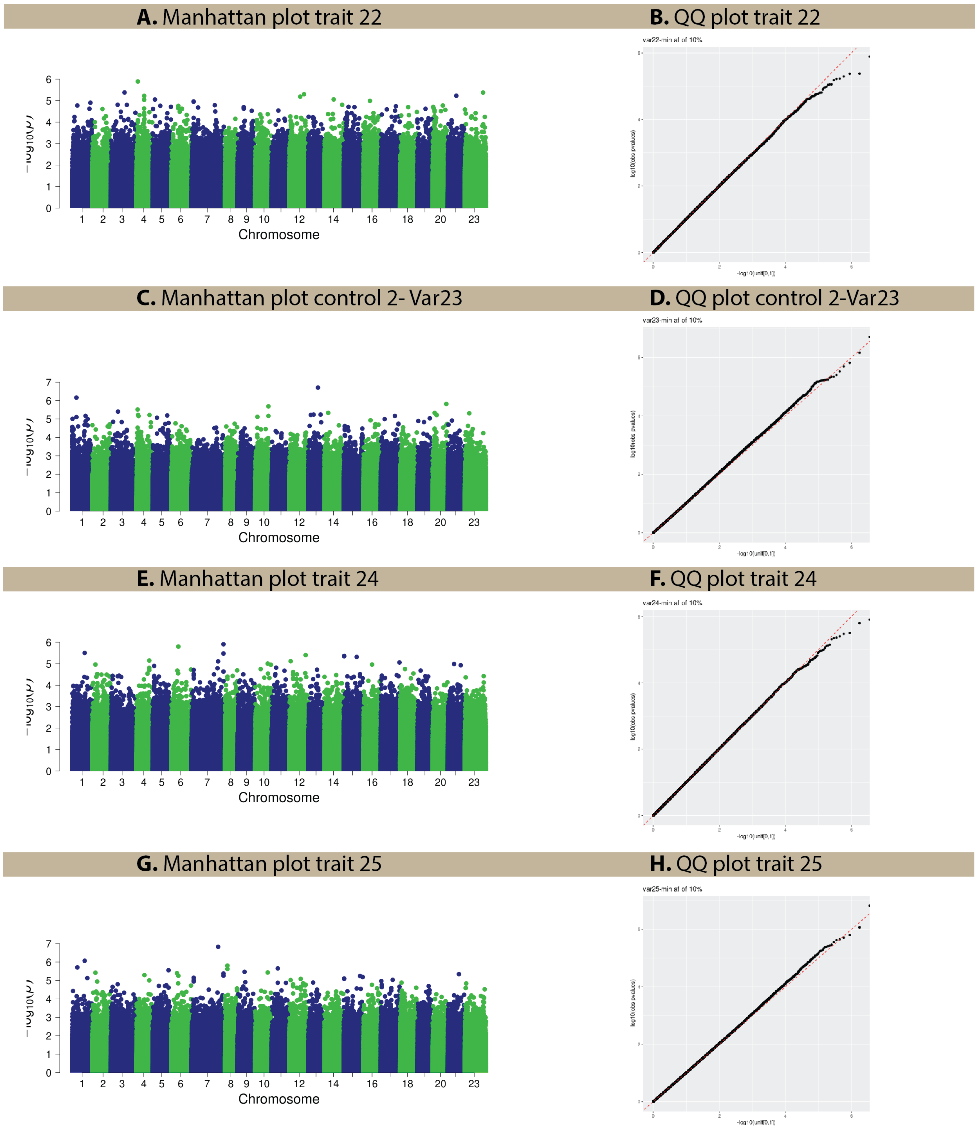


Figure Supplementary 5: (A,C,E,G) Manhattan plots for traits 22,23,24, and 25. (B,D,F,H) QQ plots for traits 22,23,24, and 25.

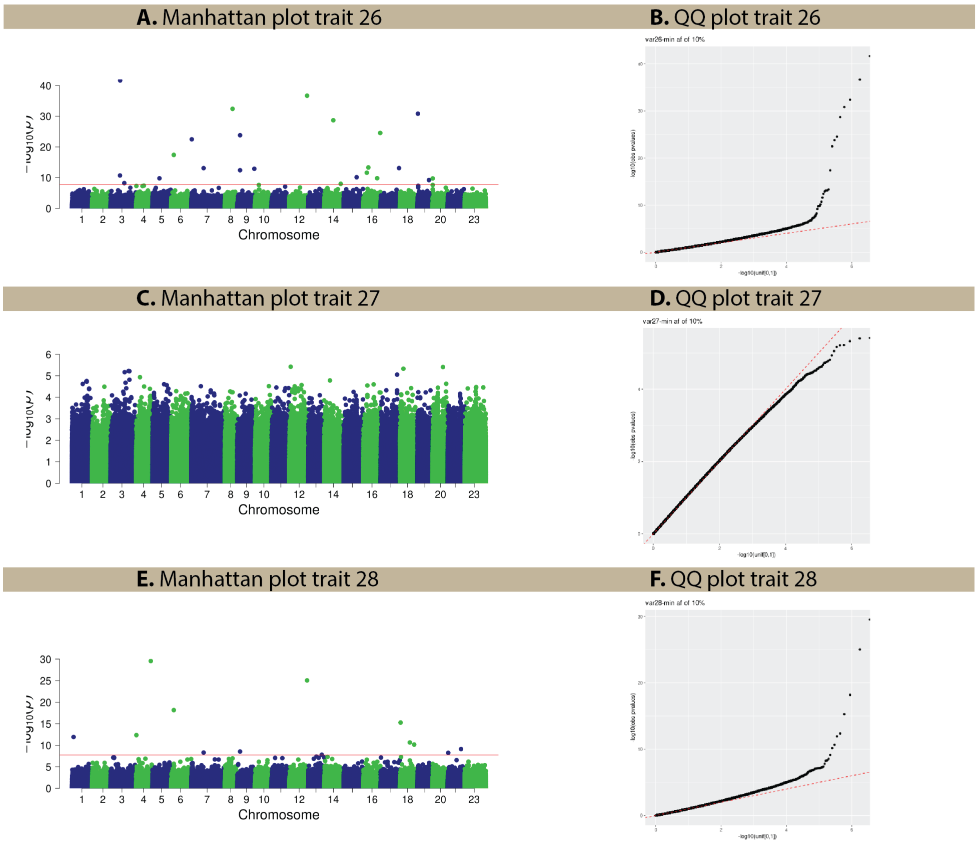


Figure Supplementary 6: (A,C,E,G) Manhattan plots for traits 26,27, and 28. (B,D,F,H) QQ plots for traits 26,27, and 28.
